# Harnessing natural diversity to identify key amino acid residues in prolidase

**DOI:** 10.1101/423475

**Authors:** Hanna Marie Schilbert, Vanessa Pellegrinelli, Sergio Rodriguez-Cuenca, Antonio Vidal-Puig, Boas Pucker

## Abstract

Prolidase (PEPD) catalyses the cleavage of dipeptides with high affinity for proline at the C-terminal end. This function is required in almost all living organisms. In order to detect strongly conserved residues in PEPD, we analysed PEPD orthologous sequences identified in data sets of animals, plants, fungi, archaea, and bacteria. Due to conservation over very long evolutionary time, conserved residues are likely to be of functional relevance. Single amino acid mutations in *PEPD* cause a disorder called prolidase deficiency and are associated with various cancer types. We provide new insights into 15 additional residues with putative roles in prolidase deficiency and cancer. Moreover, our results confirm previous reports identifying five residues involved in the binding of metal cofactors as highly conserved and enable the classification of several non-synonymous single nucleotide polymorphisms as likely pathogenic and seven as putative polymorphisms. Moreover, more than 50 novel conserved residues across species were identified. Conservation degree per residue across the animal kingdom were mapped to the human PEPD 3D structure revealing the strongest conservation close to the active site accompanied with a higher functional implication and pathogenic potential, validating the importance of a characteristic active site fold for prolidase identity.

## Introduction

Human peptidase D (PEPD) or prolidase (EC 3.4.13.9) is a multifunctional manganese-requiring homodimeric iminodipeptidase. Its enzymatic activity was reported in 1937 for the first time with the observation of Glycyl-Proline dipeptides degradation ^1^. PEPD belongs to the metalloproteinase M24 family. Its major function is the hydrolysis of peptide bonds of imidodipeptides with a C-terminal proline or hydroxyproline, thus liberating proline and hydroxyproline, respectively ^2^.

The biological significance of *PEPD* is indicated by the presence in the genomes of most animal species and its expression in several tissues ^3–7^. Moreover, *PEPD* has been identified in fungi ^8,9^, plants ^10^, archaea ^11^, and even bacteria ^12–15^. Especially the presence of PEPD in several mycoplasma species stresses its essential role in their metabolism and maintaining cellular functions, as these intracellular parasites display an otherwise extremely reduced gene set ^16^.

### Physiological role of PEPD

PEPD is the only known metalloenzyme in eukaryotes catalysing the hydrolysis of X-P ^17^. Therefore, deleterious mutations in *PEPD* in human lead to a rare autosomal disease called prolidase deficiency (PD), which is characterized by skin ulcerations due to defective wound healing, immunodeficiency, mental retardation, splenomegaly, recurrent respiratory infections and imidodipeptiduria ^18–20^. To date, 29 different pathogenic variants have been reported and associated with PD, resulting in a partial or complete enzyme inactivation ^21^. In addition to this autosomal disease, perturbations in PEPD expression, (serum) activity or serum levels have been associated with several (patho)physiological processes, including remodelling of the extracellular matrix, inflammation, carcinogenesis, angiogenesis, cell migration, and cell differentiation ^22–27^. Moreover, alterations of PEPD serum activity are associated with a spectrum of mental diseases, like post-traumatic stress disorder ^28^ and depression ^29^. Altered PEPD activity and serum level have also been frequently described in different cancer types suggesting an involvement of PEPD in cancer ^2,23,24,48^.

In bacteria and archaea, PEPD is assumed to be involved in the degradation of intracellular proteins and proline recycling ^30^. In animals, PEPD is involved in the degradation proline-rich dietary proteins and seems to play an important role in proline recycling ^2^. Since collagen (a major components of extracellular matrix) consists of 25% proline and hydroxyproline, PEPD activity is thought to be the rate limiting factor in collagen turnover ^2,31^. Interestingly, there is a growing body of evidence showing that PEPD may also have additional pleiotropic effects, independent from its enzymatic activity. Thus, PEPD has been reported to influence the p53 pathway by direct protein-protein interaction ^32^ and acts as ligand for EGFR and ErbB2 when released by injured cells ^33,34^.

### Characterization of the enzymatic and structural properties of PEPD

The crystal structure of PEPD has been extensively investigated in several species, including bacteria ^16,35^, archaea ^36^, and eukaryotes ^17^. PEPD belongs together with methionine aminopeptidase (MetAP; EC 3.4.11.18) and aminopeptidase P (APP; EC 3.4.11.9) to the “pita-bread” family, which is able to hydrolyse amido-, imido-, and amidino-containing bonds ^37,38^. Characteristic for this family is the highly conserved characteristic pita-bread fold in the catalytic C-terminal domain including the metal centre and a well-defined substrate binding pocket ^37,39^. The catalytic C-terminal domain comprises five highly conserved residues for the binding of the metal cofactors: D276, D287, H370, E412, and E452 (positions refer to human sequence) ^17^.

The preferred substrate, optimal pH and temperature, and required metal ions (e.g. Mn^2+^, Zn^2+^ or Co^2+^) are species-dependent ^2^. Although PEPD appears to be a (homo)dimer in most species including humans, it can be also active as a monomer or even as a tetramer in certain species ^2^. The homodimeric human PEPD preferably hydrolyses G-P, is adapted to a pH value of 7.8 with a temperature optimum of 50°C, and shows long-term activity at 37°C ^17,40^. *In vitro* studies based on recombinant PEPD produced in CHO cell lines and *E. coli* as well as endogenous PEPD of human fibroblasts, revealed G-P as preferred substrate followed by a lower substrate specificity for A-P, M-P, F-P, V-P, and L-P dipeptides ^40^. Moreover, in human PEPD the substrate specificity for dipeptides is determined through the presence of specific residues, like R398 and T241, which prevent the binding of longer substrates ^17^.

### Regulation of PEPD

PEPD is a phosphotyrosine and phosphothreonine/serine enzyme ^41,42^. Phosphorylation results in an increase of PEPD activity and is mediated by the MAPK pathway and NO/cGMP signalling for tyrosine and threonine/serine residues, respectively ^41,42^. Phosphorylation mediated up-regulation of PEPD activity was reported without an increased gene expression, indicating the importance of post-translational modification in its regulation ^41,42^. *In silico* analysis of human PEPD indicated post-translational modifications like glycosylations. N-glycosylation was predicted for N13 and N172, while O-glycosylation was thought to effect T458 ^22^.

We anticipate the detailed profiling of conserved residues in PEPD during evolution may help to identify and understand essential components for mentioned PEPD functions and structure. This increased knowledge could help explain the role of PEPD in diseases, especially prolidase deficiency. Taxon-specific conservation of residues provides additional insights e.g. into post-translational modification in eukaryotes. This study identified orthologous sequences of PEPD in peptide sequence sets of several hundred organisms including bacteria, archaea, animal, fungi, and plant species to investigate the conservation of residues in PEPD across the tree of life. We further identified highly conserved residues, which are likely to play key functional roles.

## Results and Discussion

### Sequence lengths differentiate between high-level taxonomic groups

In total, 769 putative PEPD orthologues were identified in animals (440), plants (122), fungi (72), archaea (42), and bacteria (93) (Supplementary File 1). PEPD orthologues in animals revealed an average sequence length of 493 amino acids (aa), while plants and fungi orthologues had an average sequence length of 499 aa and 507 aa, respectively (Supplementary File 2). Compared to these three kingdoms, PEPD sequences of bacteria were slightly smaller with an average sequence length of 455 aa. However, PEPD orthologues identified in archaea showed the smallest average sequence length of a kingdom with 360 aa. These findings matched previous reports of 349 aa (*P. furiosus*) and 493 aa (*H. sapiens*) ^11,17^. In general, our observations indicate that PEPD sequence length has changed during evolution. This length difference could be due to an increase of complexity and functionality of PEPD in eukaryotes, where it is known as a multifunctional enzyme ^2^, or due to a loss of domains in prokaryotes. Observing longer version in eukaryotes is not surprising, because eukaryotes are probably more likely to tolerate larger proteins than bacteria due to differences in the relative metabolic burden ^43^.

### Analysis of previously described residues

Our broad taxonomic sampling captured vast natural diversity, which was harnessed to identify highly conserved residues. From conservation of amino acid residues over billions of years during evolution, we infer functional relevance. A huge diversity of different species and thus sequences is key to distinguish relevant residues from the phylogenetic background. To ensure an accurate alignment of all analysed sequences, the alignment was performed with permutations of the input sequences and repeated with different alignment tools. The average difference per position in the resulting alignments is low (Supplementary File 3 and 4).

### Conservation of functional and structural relevant residues

Highly conserved residues are likely to have a high functional, and/or structural relevance. Aiming to extend the knowledge about the already existing crystal structure of especially human PEPD, we analysed the conservation degree of known residues relevant for the structure and function of PEPD ^17^. Despite the high diversity of metal ions accepted by different species ^2^, the amino acids responsible for the binding of the metal ions (D276, D287, H370, E412, and E452) are highly conserved across species (Supplementary File 5). All residues reported for the interaction with metal ions were detected in over 90% of all sequences. Sequences without these particular residues are likely to be partial and thus not covering this position leading to a lower observed conservation value. When excluding sequence gaps, almost 100% match is reached for all five positions. Based on these results, we conclude that all selected sequences are *bona fide* prolidases. This finding marks the conservation of these five residues as one important structural and functional characteristic of PEPD (Figure 1).

**Figure 1:**
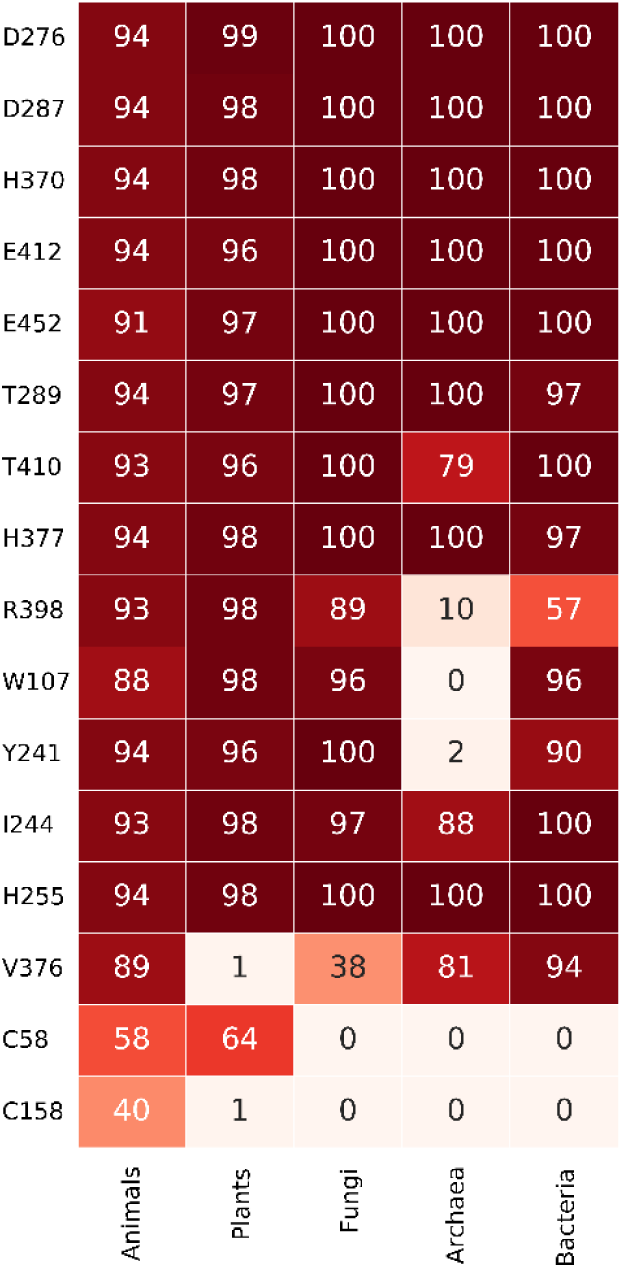
Heatmap of reported functionally important residues of PEPD. The conservation degree of reported residues important for PEPD functionality and structure is displayed in percentage across species. Each column represents a kingdom, while the rows display the analysed residue and its corresponding position in the human PEPD amino acid sequence. A dark green background indicates high conservation, while white means no conservation.

Additionally, strong conservation of T289 and T410 in proximity to the manganese ions supports previous reports and hypotheses of their functional relevance in PEPD ^22^.

Nevertheless, one plant- and three animal PEPD orthologues showed an amino acid substitution of one metal binding residue: *Ancylostoma ceylanicum* (H370V), *Arachis duranensis* (D287N), *Oncorhynchus kisutch* (E452K) and *Tetraodon nigroviridis* (E452R). Crystal structures and enzyme assays could illuminate the consequences of these substitutions thus providing natural sequences to assess the contribution of each residue. Since D287N was reported before as a probably deleterious substitution ^44^, these prolidases may have lost their ability to cleave X-P dipeptides.

Another essential step for the enzymatic catalysis of prolidases is the binding of their dipeptide substrate (e.g. G-P)^17^. For example, H255 binds to the carboxylate group of the C-terminal proline residue of the substrate and its side chain moves upon substrate binding by about 6 A° narrowing down the size of the active site ^17^. The importance of such substrate binding residues, like H255 and H377 ^17^, was validated through a high conservation degree of minimum 94% in all living organisms (Figure 1). Interestingly, another residue involved in G-P binding in human PEPD, R398 ^17^, is highly conserved except in archaea (Figure 1). Besides its role in G-P binding, this residue is also important for the specificity of PEPD for dipeptides by determining the length of the ligand at the C-terminus through its large side chain ^16,17^. These results suggest that the majority of analysed archaeal prolidases might not be capable of G-P degradation and may have a broader substrate spectrum due to the missing R398. In line with the hypotheses, Ghosh *et al.* showed that PEPD purified from the archaeon *P. furiosus* revealed no substrate specificity for G-P, but for longer substrates like K-W-A-P and P-P-G-F-S-P, although this specificity was rather weak ^11^. However, the preferred substrates of this enzyme were the dipeptides M-P and L-P ^11^. Interestingly, *P. furiosus* still has a corresponding arginine residue at the position 295 ^16^. This R295 was reported to have dual functionality for cleaving di- and tripeptides due to the intermediate position of this arginine ^16^. These reports support the hypothesis that archaeal prolidases have a broader substrate spectrum compared to the prolidases of the other kingdoms. In turn, the strong conservation of R398 in eukaryotes may indicate an adaptation to the specific recognition of dipeptides. In line with the hypothesis, the bulky side chain of R398 was reported to prevent the acceptance of tripeptides ^17^. Moreover, a strong conservation of W107, except in archaea, was identified (Figure 1). After G-P binding, W107 is shifted inwards to the active site, sealing the active site ^17^. The low conservation of W107 in archaea suggests that archaeal prolidases might use a different conformational change, probably due to their putative expanded substrate spectrum.

Furthermore, some residues were reported to be involved in the interaction of L-P, another potential prolidase substrate: Y241, I244, H255, and V376 ^17^. H255 and I244 are highly conserved across species (Figure 1). V376 is less conserved in fungi and not conserved in plants. Y241 is not conserved in archaea. Since *P. furiosus* PEPD is capable of binding and degrading L-P, Y241 is probably not essential for this binding process in archaea. Another reason for the flexibility in archaea might be the putatively expanded substrate spectrum due to the absence of Y241, which is reported to close the active site on the side where the N-terminus of the substrate is placed ^21^. To the best of our knowledge, the effect of the absence of V376 in plants was not investigated yet.

In order to identify a common disulfide bond responsible for the common dimer formation of prolidases previously reported cysteine residues ^17^ were analysed. In human PEPD an intramolecular disulfide bridge was observed between C58 from chain A and C158 from chain B ^17^. However, this bond was only present in the inactive (Mn^2+^ free) enzyme complex, while the substrate was bound in the active site ^17^. These amino acids are weakly conserved in the animal kingdom (58% and 40% respectively), but showed an almost complete conservation among vertebrata likely due to their relevance in the dimer formation in this group. However, these cysteines might not be responsible for the dimer formation in the active form of the enzyme, which occurs in most of the prolidases ^8,17,45^. Therefore, we aimed to identify a better candidate for this common PEPD conformation. However, we could not identify a highly conserved cysteine across species, suggesting (I) the presence of different interactions for stabilization of e.g. PEPD dimers or (II) frequent occurrence of PEPD as a monomer.

### Analysis of residues known to be mutated in prolidase deficiency

The majority of amino acids that are hot spots causing PD (6/11: D276, G278, L368, E412, G448, G452) are localised near or in the active side of PEPD ^22,46^. These amino acids are conserved across species, thus suggesting a negative correlation between the distance of a residue to the active site and its conservation in animals. As expected, highly conserved (>85%) residues are more likely to be located close to the active site (p-value= 3.76e-06, Mann-Whitney U test)(Figure 2, Supplementary File 6).

**Figure 2:**
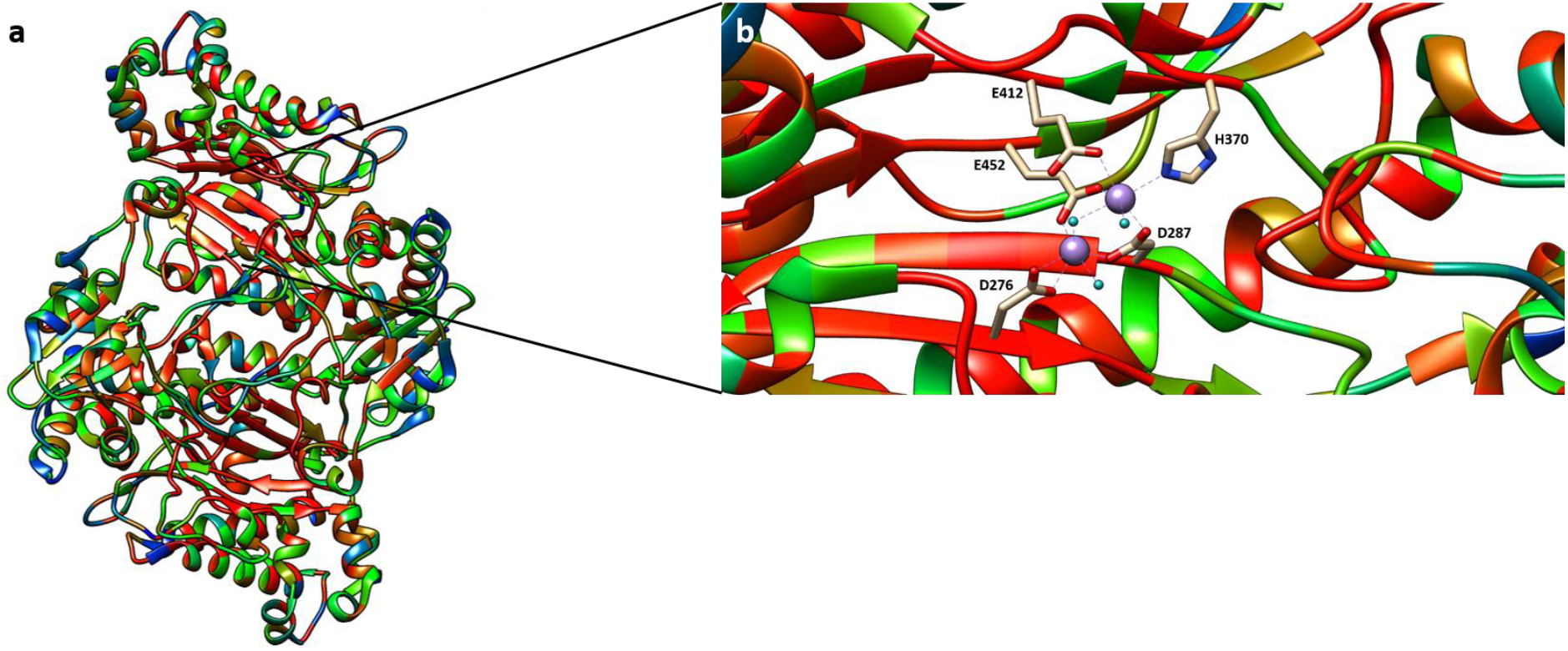
The catalytic cavity is highly conserved in the animal kingdom. (a) Three dimensional heatmap of residue conservation degree in the animal kingdom, represented by the PEPD structure of human prolidase (5M4G). The colour scale ranges from red (highly conserved residues) over orange and green to blue (weakly conserved residues). (b) Conservation degree of the catalytic site of human PEPD. The metal binding residues (D276, D287, H370, E412, and E452) are shown together with the bound Mn^2+^ ions (violet) and water molecules (cyan).

As mentioned previously the metal binding residue E452 is highly conserved across species and its deletion results surprisingly in a preservation of the active site ^21^, likely because it can be replaced by neighbouring residues. However, the mutated protein shows less than 5% of the WT activity ^47^ supporting our findings. Additionally, our results are in line with findings of Bhatnager and Dang, who identified the mutation of D276N, G278D, E412K, and G448R as damaging substitutions ^44^, because we observed a strong conservation of all four residues. Recently the structural basis of these and other PD mutations have been analysed in detail ^21^. Once again in accordance with our results, Wilk *et al.* claimed that the D276N mutation results in an excessive reduction of the PEPD activity due to the loss of one of the catalytic metal ions derived from the charge change caused by the substitution ^21^. Similarly, in the G278D mutant the loss of one metal ion and additional enhanced disorder were observed ^21^. Interestingly, the previously as highly conserved identified Y241 seems to have high functional relevance since its displacement in this mutant results in a destabilization of two metal binding residues (D276 and D287)^21^. In addition, the highly conserved substrate coordinating residue H255 is completely absent from the active site of the G278D mutant ^21^ stressing its importance in maintaining PEPD functionality. H255 is also absent in the G448R mutant contributing to a dysfunctional protein core ^21^. The substitution of the metal binding E412 to K results once again in the loss of one metal ion by an amino acid side chain leading to PEPD inactivation ^21^.

R184 is defined by the shortest atom-to-atom distance to G-P in human PEPD and marks the end of the N-terminal chain of human PEPD ^21^. The deletion or mutation of R184 to Q in PD patients results in an inactive PEPD or one with highly reduced enzyme activity, respectively ^21^. Therefore, R184 might be essential for the functionality and structure of PEPD, which is supported by its high conservation across many species ^22^. In this study, this finding was validated with a minimum conservation degree of 92% of all sequences analysed. Moreover, D375 and D378 were identified as highly conserved across species. Interestingly, these residues were both recently reported to directly interact with R184 ^21^. In the PD mutation variant R184Q, the interaction between R184 with D375 and D378 is lost, due to the replacement of the positive charged guanidinum group of R184 to the neutral amide group of Q ^21^. The resulting protein shows only residual activity, supporting the hypothesis that D375 and D378 are highly important for PEPD functionality.

Additional relevant residues in PD are not particular conserved across different phyla. Among them are S202 (90%) and Y231 (89%) highly conserved in animals. While the deletion of Y231 results in alterations in the dimer interface with remaining PEPD activity, the S202F substitution increases PEPD disorder resulting in the inability to hydrolyse G-P ^21^. Y241 is affected by S202F contributing to loss of PEPD activity, since Y becomes disordered even though all other metal binding residue are not affected ^21^. Since Y241 interacts in the WT human PEPD structure with the metal binding aspartates ^21^, its disorder might result in the loss of this interaction, thus destabilizing PEPD. However, A212 (45%) and R265 (35%) show a substantially smaller conservation degree compared to S202 and Y231. Strong conservation of A212 and R265 is limited to vertebrates thus suggesting a pathogenic role limited to this branch. The phenotype of S202P, A212P, and L368R are not distinguishable from each other, posing an example for relevant residues in PD without strong conservation ^46^.

### Identification of polymorphisms in damage-associated SNPs in human prolidase gene

Recently, Bhatnager and Dang (2018), identified damage associated single-nucleotide polymorphisms (SNPs) in human prolidase gene based on a comprehensive *in silico* analysis ^44^. We observed that some of their non-synonymous SNPs are leading to substitutions at variable positions thus qualifying as polymorphisms instead of pathogenic variants. Such a SNP is causing the substitution of V to I at position 305, while our analysis revealed V in 78% and I in 16% of all animal PEPD sequences. Six out of seven tools predicted this SNP as neutral, supporting our assumption ^44^. Similar ratios and even dominance of a different amino acid were observed for I45V, E227L, and L435F indicating three additional polymorphisms. Additionally, we hypothesize that nsSNPs leading to T137M, V456M, and D125N are likely to be polymorphisms as the conservation of the canonical amino acid is low.

However, the remaining nsSNPs showing a higher conservation degree in the animal kingdom indicate that they may be important for structure or function of PEPD in the animal kingdom and that substitutions of these residues have a pathogenic potential ^44^. This is especially the case for the overlaps of the identified consensus nsSNPs, which were predicted from all tools as damage associated, with our results stressing that these residues are highly conserved not only in the animal kingdom, but also across species ^44^(Table 1).

**Table 1:** Conservation degree across species for positions, which were reported to be derived from damage-associated nsSNPs. The conservation degree of positions, which were reported to be derived from damage-associated nsSNPs are stated for animals (An), plants (Pl), fungi (Fu), bacteria (Ba) and archaea (Ar). The first column contains the position of each amino acid based on the human PEPD sequence (Reference sequence position, RSP; UniProt ID: P12955). The amino acid frequency (AAF) ranging from 0 to 1 (1=100% conserved) of the most abundant (1) and second abundant (2) amino acid at a certain position is listed. Gaps in the alignment are indicated through a “-“ followed by the conservation degree in the kingdom. Only a “-“ is given, when the first amino acid is 100% conserved.

| RSP | An AAF1 | An AAF2 | Pl AAF1 | Pl AAF2 | Fu AAF1 | Fu AAF2 | Ba AAF1 | Ba AAF2 | Ar AAF1 | Ar AAF2 |
| --- | --- | --- | --- | --- | --- | --- | --- | --- | --- | --- |
| 19 | P_0.71 | S_0.18 | P_0.73 | -_0.1 | P_0.97 | D_0.01 | -_0.62 | P_0.26 | -_1.0 | - |
| 35 | R_0.51 | K_0.24 | R_0.76 | -_0.06 | L_0.31 | R_0.17 | P_0.37 | A_0.22 | E_0.25 | Y_0.1 |
| 188 | T_0.73 | S_0.2 | S_0.89 | T_0.08 | D_0.82 | T_0.1 | D_0.5 | T_0.43 | D_0.63 | E_0.13 |
| 192 | L_0.67 | I_0.25 | L_0.8 | I_0.14 | I_0.64 | V_0.19 | I_0.38 | L_0.34 | I_0.5 | L_0.38 |
| 224 | S_0.81 | A_0.11 | S_0.95 | A_0.02 | A_0.92 | G_0.06 | G_0.28 | L_0.25 | A_0.43 | G_0.2 |
| 240 | S_0.84 | A_0.08 | S_0.90 | -_0.02 | A_0.38 | G_0.35 | P_0.44 | G_0.4 | S_0.48 | A_0.48 |
| 247 | S_0.79 | T_0.13 | T_0.89 | S_0.07 | S_0.74 | A_0.21 | L_0.41 | S_0.24 | S_0.45 | F_0.38 |
| 255 | H_0.94 | -_0.05 | H_0.98 | -_0.02 | H_1.0 | - | H_1.0 | - | H_1.0 | - |
| 276 | D_0.94 | -_0.05 | D_0.99 | -_0.01 | D_1.0 | - | D_1.0 | - | D_1.0 | - |
| 278 | G_0.94 | -_0.06 | G_0.99 | -_0.01 | G_0.97 | A_0.03 | G_0.99 | T_0.01 | G_0.95 | T_0.05 |
| 287 | D_0.94 | -_0.05 | D_0.98 | -_0.02 | D_1.0 | - | D_1.0 | - | D_1.0 | - |
| 296 | G_0.94 | -_0.05 | G_0.98 | -_0.02 | G_0.97 | T_0.01 | G_0.62 | S_0.19 | G_0.45 | -_0.18 |
| 373 | G_0.94 | -_0.06 | G_0.98 | -_0.02 | G_1.0 | - | G_1.0 | - | G_1.0 | - |
| 378 | D_0.93 | -_0.05 | D_0.98 | -_0.02 | D_1.0 | - | D_0.94 | E_0.06 | E_0.85 | D_0.15 |
| 403 | L_0.80 | V_0.12 | L_0.96 | -_0.02 | L_0.94 | V_0.04 | L_0.78 | I_0.15 | L_0.85 | I_0.13 |
| 410 | T_0.93 | -_0.06 | T_0.96 | -_0.02 | T_1.0 | - | T_1.0 | - | T_0.78 | S_0.23 |
| 412 | E_0.94 | -_0.06 | E_0.96 | -_0.02 | E_1.0 | - | E_1.0 | - | E_1.0 | - |
| 447 | G_0.93 | -_0.07 | G_0.97 | -_0.03 | G_1.0 | - | G_0.53 | -_0.4 | F_0.6 | G_0.25 |
| 448 | G_0.93 | -_0.06 | G_0.97 | -_0.03 | G_1.0 | - | G_1.0 | - | G_1.0 | - |

### PEPD in cancer

The investigation of curated SNPs in *PEPD*, which are associated with specific cancer types (BioMuta database ^49^), revealed missense mutations in various cancer types to be distributed across the whole PEPD sequence (Supplementary File 7). As many SNPs were associated with a low frequency, we focused on a small set of more frequent ones. Surprisingly, the amino acid affected by the most frequent SNPs in various cancer types is A74, a residue located in the non-catalytic N-terminal domain. While the general frequency in animals is low (38%), it displays a strong conservation in mammals thus suggesting a functional role. Other frequently effected residues are A122, H155, G257, R311, M329, and D378. All of them are conserved to different extents in the animal kingdom, while three (G257, M329, and D378) are also conserved in plants. However, D378 is the only amino acid conserved across all species. Being in proximity to the metal binding residue H370, the high conservation degree of D378 might be due to its role in forming a functional catalytic site. However, we could not identify a “cancer specific hot spot residue” in the animal kingdom and thus the appearance of SNPs in *PEPD* in various cancer types is likely not to be the driving force of a specific cancer type and the identified SNPs might be polymorphisms.

### Post-translational regulation of PEPD

Since there is experimental evidence of PEPD activity being regulated at the post-translational level through phosphorylation ^41,42^, we aimed to validate previously predicted post-translational modifications (PTMs) ^50^ in human PEPD. None of the examined sites were highly conserved across species (Supplementary File 5), which could be explained by differences in the PTM mechanisms between prokaryotes and eukaryotes ^51,52^. Nevertheless, some residues were conserved in the animal kingdom e.g. R196 (88%). The low conservation values could be due to differences in PTMs between different groups of eukaryotes ^51^. The lack of conservation for some of these residues (S8, K36, S113, T487, A490, K493) could be explained in three ways: (I) no strong functional relevance for PEPD, (II) false positive prediction, or (III) a human specific regulation system. *Vice versa*, three residues are highly conserved at least in the animal kingdom (T15:80%, Y128:78%, R196:88%) posing good candidates for a PTM site. Two of the three amino acids are predicted to be phosphorylated (T15 and Y128), while R196 is thought to be monomethylated ^50^.

Lupi *et al.* predicted putative PTMs at N13, N172 (NetNGly), and T458 (NetOGlyc) ^22^. These residues were found to be highly conserved among vertebrates. This situation could be explained by a more recently evolved function or a relaxed ancestral function in species without strong conservation. *In silico* prediction of new phosphorylation sites resulted in T90, S113, Y121, Y128, S202, S224, S138, S240, S247 and S460 as best candidates. Conservation degrees generally support these predictions (Supplementary File 5) and distribution across species suggests a more recently increased relevance of S113 and S138.

### Identification of novel conserved residues

All structure related observation and hypothesis are based on human prolidase crystal structure (PDB: 5M4G). As we already validated through the correlation in the animal kingdom, highly conserved residues are located nearby or in the substrate binding site. Therefore, it was not surprising that residues near the metal binding residue E452 are highly conserved across species especially R450:92% along with the previously reported G448:93%. The side chain of R450 is near the metal binding site, indicating that it might be essential for the formation of a functional metal ion binding site (Supplementary File 8 (a)). Another two conserved residues, T458 and G461, are located in the curve of a C-terminal loop near the binding site (Supplementary File 8 (b)). The small size of these amino acids might be necessary to form this structural feature. However, T458 could be a putative phosphorylation site. Since it is located on the outer surface of the enzyme, it is accessible for modifications. Additionally, we observed a cluster of highly conserved residues (G406-V408), which are part of the pita-bread structure, stressing the importance of this fold for the function of PEPD as metalloproteinase.

Again, highly conserved residues across species were identified near another known metal binding residue E412: Y416:94%, P413:94%, and G414:93% are located near the active site and are therefore good candidates for generating a functional binding site. The glycine and proline seem to be important to allow the proper arrangement of the metal binding residues by providing space between them. The side chain of Y416 is pointing into the active side, indicating it might have an additional functional role (Figure 3 (a)).

**Figure 3:**
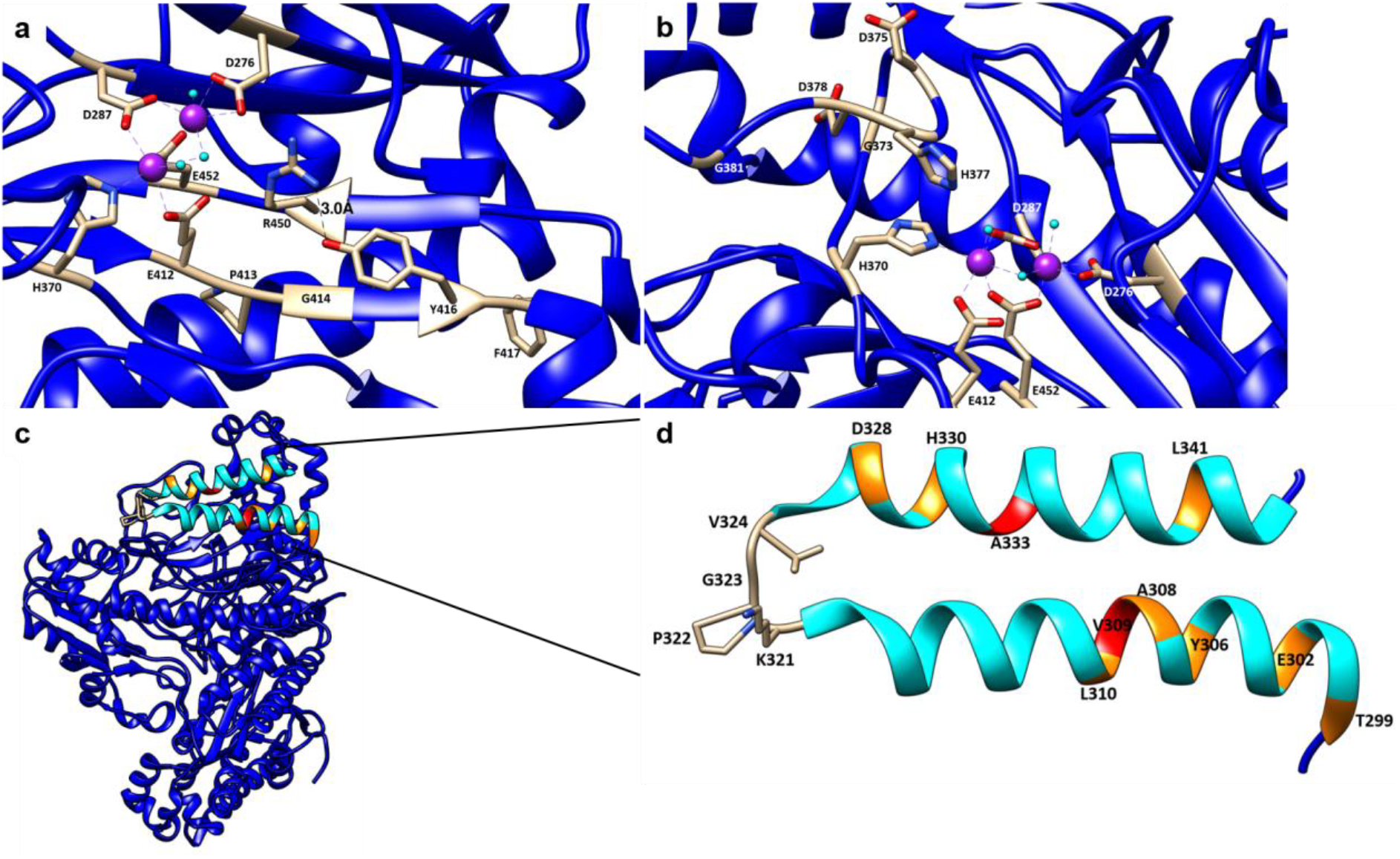
Novel highly conserved residues with functional and/or structural importance in PEPD. The ribbon of the human PEPD 3D model is shown in blue, while residues of interest are lettered. The metal ions are shown in violet and water molecules are shown in cyan. (a) Highly conserved residues P413, G414, and Y416 are located near the metal binding residue E412 and are likely to be involved in generating a functional binding cavity. Y416 might stabilize the anti-parallel β-strand through interaction with R450. (b) G373, D375, D378, and G381 are involved in the stabilization of the loop, which results in an optimal position of the substrate-binding residue H377. (c) Peripheral localisation of the helix with highly conserved residues. (d) The peripheral helix contains two highly conserved residues (A333 and V309), which are marked in red and other conserved residues, which are marked in orange. Moreover, residues building the loop (V324, G323, P322, and K321) are conserved, too.

However, it is more likely that it has a stabilizing effect building a hydrogen bond with the NH group of R450:92% (Figure 3 (a)) thus stabilizing the anti-parallel β-strand. This anti-parallel β-strand seems to be highly important for PEPD functionality, since substitutions in the parallel β-strand e.g. G447R or G448R were reported to null PEPD activity ^44^. The insertion of a bulky arginine side chain, which prevents the correct assembly of the β-sheet, could be the explanation ^44^. Furthermore, F417:82% is highly conserved in every kingdom except archaea, expanding the number of conserved residues in this conserved region (Figure 3 (a)).

The conserved G373 is located in a tied turn of the peptide chain, suggesting its interplay with the conserved residues D375, D378, and G381 to form a loop. As a result, the important dipeptide-binding residue H377 is placed near the catalytic site (Figure 3 (b)). Weak conservation of these residues in archaea vindicates the previously mentioned hypothesis that archaea PEPD might be able to hydrolyze a broader substrate spectrum. Additionally, we identified the two conserved residues G369 and H366 near the metal binding residue H370 (Supplementary File 8 (c)). The side chain of H366 is pointing into the active site, indicating that it will narrow down the active site, therefore contributing to substrate specificity. Interestingly, residues near H366 e.g. P365, G367, and L368 are highly conserved with exception of the archaea kingdom. This could explain the ability of archaeal prolidases to process tripeptides in addition to dipeptides.

The highly conserved residues T299, E302, Y306, A308, V309, L310, K321, P322, G323, V324, D328, H330, and L341 form two parallel helices located in the periphery of PEPD, thus exposed to the solvent (Figure 3 (c)). Based on their extremely high conservation, V309 and A333 are probably most important for this structure (Figure 3 (d)). Whether this region could be the cause for some of extracellular functions of PEPD, e.g. EGFR or ErbB2 binding ^33,34^ or might be a target for a regulatory protein, needs to be investigated in the future.

Moreover, T299, F298, G296, and P293 are highly conserved across species except archaea. These residues might stabilize the pita-bread fold by strengthening a loop near the catalytic site (Supplementary File 8 (d)). Additionally, near the metal binding residue D276, some amino acids display strong conservation including G278, G270, E280, and L274.

Interestingly, investigation of residues near the highly conserved H255 revealed an exclusive conservation of the region between L257 and A259 in animals and plants. It is located in a loop structure at the periphery of PEPD. This region and other similar observations e.g. G385, V386, M236, G149, N151, T152, Q49, and G50 indicating that plant and animal prolidases might have distinct structural features compared to archaea, bacteria, and fungi. However, the flanking amino acids of H255 are highly conserved at a minimum of 94% in animals, plants and fungi, stressing its importance in eukaryotes.

Overall, we observe more conserved residues in the C-terminal catalytic region compared to the N-terminal region. Nevertheless, P98, L95, P80, G76, and F65 are examples for conserved residues in the N-terminal part. Their functions are yet to be determined.

### Limitations and perspectives

Numerous PEPD orthologues were identified across all living organisms to pinpoint key residues in this protein. The selection of sequences from different groups is not balanced and we do not attempt to assign evolution events to certain groups, which would be possible based on an even more comprehensive sample. A high natural diversity allowed us to distinguish between variable positions with low if any functional relevance and highly conserved residues, which are likely to play key catalytic, structural, or regulatory roles in PEPD. The results match previously reported residues and enabled us to identify additional residues, which should be subjected to in-depth investigation and will eventually shed light on function and structure of PEPD. However, 264 (27%) of the screened data sets did not reveal a PEPD candidate based on our bait sequences. A majority of species without PEPD candidates (175) were bacteria (Supplementary File 9). Since PEPD is a relevant enzyme at least in eukaryotes, it is unlikely to be missing in many species. Technical limitations like incomplete assemblies or annotations could be the reasons for the absence of PEPD from some data sets. Therefore, we checked the completeness of all analysed data sets through the identification of suitable benchmarking genes that are assumed to be present in the respective species (Supplementary File 9) and discussed it in detail (Supplementary File 10). The identification of additional PEPD orthologues would facilitate further analyses e.g. improve the differentiation between pathogenic substitutions and harmless polymorphisms. We used our observations to predict the functional impact of nsSNPs and expect that this approach will be useful in the future for similar applications. We anticipate that the use of *in silico* tools integrating evolutionary genetics and structural data available will help to gain knowledge e.g. regarding the molecular characterization of PEPD, the identification of new regulatory residues, the extracellular role of PEPD, and new therapeutic strategies against prolidase deficiency and other PEPD associated disorders.

## Material and methods

### Data set collection

The peptide sequence sets of 475 animals, 122 plants, 72 fungi, 49 archaea, and 236 bacteria were retrieved from the NCBI. All sequences were pre-processed with a dedicated Python script to generate customized data files mainly with adjusted sequence names as long sequence names can pose a problem to some alignment tools (https://github.com/bpucker/PEPD). Next, peptide sequence sets were subjected to BUSCO v3 ^53^ to assess their completeness based on the reference sequence sets ‘metazoa odb9’ (animals), ‘embryophyta odb9’ and ‘eukaryota odb9’ (plants), ‘eukaryote odb9’ (fungi), and ‘bacteria odb9’ (bacteria). Since there is no dedicated reference sequence set available for archaea, we used the eukaryota and bacteria sets. PEPD bait sequences (Supplementary File 11 and 12) were selected manually based on the literature and/or curated UniProt entries ^8,36^. Initial selection of related sequences was based on a pipeline combining previously published scripts and using their default parameters ^54^. Candidate sequences were identified in a sensitive similarity search by SWIPE v2.0.12 ^55^ and filtered through iterative steps of phylogenetic analyses involving MAFFT v7.299b ^56^, phyx ^57^, and FastTree v2.1.10 ^58^. Results were manually inspected and polished to identify *bona fide* orthologous genes with a high confidence. As the average length of PEPD in animals and plants is around 500 amino acids, sequences outside the range 200-700 amino acids were filtered out to avoid bias in downstream analyses through partial sequences or likely annotation artefacts.

### Identification and investigation of conserved residues

MAFFT v.7.299b ^56^ was applied for the generation of multiple sequence alignments. Resulting alignments were cleaned by removal of all alignment columns with less than 30% occupancy. Conserved residues were identified and listed based on positions in the human PEPD sequence (UniProt ID: P12955) using the Python script ‘conservation_per_pos.py’ (Supplementary File 1). This analysis was repeated 50 times with randomly reshuffled sequences as the order of sequences can heavily impact the alignment process ^59^. In addition, we compared the alignments generated by MAFFT v.7.299b to ClustalO v.1.2.4 ^60^ and MUSCLE v.3.8.31 ^61^ alignments of the same data sets. The alignment bias through the order of input sequences was quantified for all positions of the aligned *Homo sapiens* sequence. For the *in silico* prediction of phosphorylation sites the *H. sapiens* PEPD sequence (UniProt ID: P12955) was submitted to NetPhos 3.1 ^62,63^. Only the best prediction for each residue with a high confident score of >0.8 was considered for further analyses.

### Sources of previously reported data

Previously reported residues with functional implications (Supplementary File 7) were checked for conservation. Additionally, the alignment was screened for highly conserved residues to the best of our knowledge not previously reported in respect to functionality or structure of PEPD. The results of the residue conservation analysis for the animal kingdom were mapped to a 3D structure of human PEPD (PDB: 5G4M). Putative post-translational modification sites were obtained from PhosphoSitePlus and literature ^22,50^. Residues associated with PD were retrieved from literature ^22,46^. Non-synonymous single-nucleotide polymorphisms (nsSNPs) ^44^ and details about observations were retrieved from the curated BioMuta database ^49^.

### Correlation analysis of conservation degree and distance to the active site of PEPD

To determine the conservation degree in correlation to the distance to the active site, the average localisation of the five metal binding residues was identified and used to calculate the distance of each residue to this focus of the catalytic site (Supplementary File 13). Information about the position of each residue was taken from the PDB file 5M4G of human PEPD ^17^. The Python modules matplotlib ^64^ and seaborn (https://github.com/mwaskom/seaborn) were applied to construct a conservation heatmap. In addition, the conservation of all residues in animals was mapped to the 3D model of the human PEPD by assigning colours within a colour gradient to each amino acid representing its conservation among animal sequences.

### Phylogenetic analysis

A phylogenetic tree was constructed via FastTree v.2.1.10 ^58^ based on alignments generated via MAFFT v.7.299b ^56^ and trimmed via pxclsq ^57^ to a minimal occupancy of 60%. The conservation of different key residues was mapped to this tree for visualization. A Python script (https://github.com/bpucker/PEPD) was deployed to colour all leaves representing sequences with the conserved residue in red.

## Data Availability

All data generated or analysed during this study are included in this published article (and its Supplementary Information files).

## Acknowledgements

We thank Samuel F. Brockington, Nathanael Walker-Hale, and Kali Swichtenberg for critical reading of the manuscript and very helpful comments.

## Authors’ contributions

HMS and BP designed the experiments, performed bioinformatics analyses, interpreted the results, and wrote the manuscript. All authors revised the manuscript. We acknowledge support for the Article Processing Charge by the Open Access Publication Fund of Bielefeld University.

## Supplementary material

**Supplementary File 1: PEPD peptide sequences used for multiple sequence alignments.**

**Supplementary File 2: Length distribution of PEPD orthologues.** PEPD sequence length is displayed on the x-axis, while the frequency of a sequence length in percentage is shown on the y-axis. Archaea orthologues are coloured in violet, bacteria in black, fungi in blue, plants in green, and animals in red.

**Supplementary File 3: Alignment bias control.** The y-axis displays the conservation degree ratio of each residue across species as well as the variation of this value between alignments (Supplementary File 4). The x-axis shows the corresponding residue position in the human PEPD amino acid sequence (UniProt ID: P12955). The green line shows the median of all conservation values observed across all generated alignments. The red line displays the maximum conservation degree and the blue line the minimum conservation degree observed for the respective position across all alignments, respectively.

**Supplementary File 4: Alignment bias control values.** The variation of the calculated conservation degree based on multiple alignments by MAFFT, ClustalO, and MUSCLE is listed. The first column contains the position in the reference sequence human PEPD (UniProt ID: P12955). In addition, the minimal conservation degree observed over 50 alignments, the median of all these conservation values, and the maximal observed value are provided.

**Supplementary File 5: Conservation degree of PEPD residues across species.** The conservation degree of each residue, ranging from 0-1.0 (1.0 being perfect conservation) is listed for animals, plants, fungi, bacteria, and archaea. The alignment position of each residue is given in the first column, while the second column refers to the corresponding position in human PEPD (Reference sequence position, UniProt ID: P12955). The amino acid frequency (AAF) of the most abundant (AAF1) and second abundant amino acid (AAF2) at a certain position is given for each species. A gap is indicated by “-“.

**Supplementary File 6: Distance of each residue to the active site of human PEPD.** The distance of each residue to the active site of human PEPD (PDB ID: 5M4G) is stated in arbitrary units.

**Supplementary File 7: Previously reported residues for conservation analysis.** All previously reported residues with relevance to structure and/or function of PEPD are listed with their associated function and reference. The residue position is derived from human PEPD (UniProt ID: P12955). PTMs identified in *H. sapiens* or *M. musculus* are marked through Hs and Mm in brackets, respectively.

**Supplementary File 8: Conserved residues in human PEPD 3D model with structural and/or functional relevance.** The ribbon of the human 3D PEPD model is shown in blue, while residues of interest are marked in red or alternatively in beige. The metal ions are displayed in violet and water molecules are shown in cyan. (a) R450 (highlighted in red) is located near the metal binding centre. (b) T458 and G461 are marked in red and are located in a peripheral loop. (c) G369 and H366 are located near the metal binding residue H370, where H366 might narrow down the active site. Moreover, P365, G367, and L368 might be involved in substrate specificity of animal, plant, fungi, and bacteria PEPD.

(d) T299, F298, G296, and P293 stabilize the pita-bread fold by strengthening the loop near the catalytic site.

**Supplementary File 9: BUSCO assessment of peptide data set quality.** For each analysed organism presence (+) or absence (-) of PEPD in their peptide dataset is indicated. Completeness of the data sets was assessed based on the detection of BUSCO sequences.

**Supplementary File 10: Discussion of possible limitations.**

**Supplementary File 11: Identifier of bait sequences.** Donor species and NCBI or UniProt ID of PEPD bait sequences is listed.

**Supplementary File 12: Bait sequences.**

**Supplementary File 13: Approach for residue distance calculation.** Schematic illustration of the approach used to calculate the distances of all amino acids in PEPD to the active site. Different colours indicate different amino acids with different degrees of conservation across species.

## References

1. Bergmann, M. & Fruton, J. On proteolytic enzymes. XII. Regarding the specificity of aminopeptidases and carboxypeptidases. A new type of enzyme in the intestinal tract. J. Biol. Chem. 177, 189–202 (1937).

2. Kitchener, R. I. & Grunden, A. m. Prolidase function in proline metabolism and its medical and biotechnological applications. J. Appl. Microbiol. 113, 233–247 (2012).

3. Davis, N. C. & Smith, E. L. Purification and some properties of prolidase of swine kidney. J. Biol. Chem. 224, 261–275 (1957).

4. Baksi, K. & Radhakrishnan, A. N. Purification and properties of prolidase (imidodipeptidase) from monkey small intestine. Indian J. Biochem. Biophys. 11, 7–11 (1974).

5. Browne, P. & O’Cuinn, G. The purification and characterization of a proline dipeptidase from guinea pig brain. J. Biol. Chem. 258, 6147–6154 (1983).

6. Endo, F., Hata, A., Indo, Y., Motohara, K. & Matsuda, I. Immunochemical analysis of prolidase deficiency and molecular cloning of cDNA for prolidase of human liver. J. Inherit. Metab. Dis. 10, 305–307 (1987).

7. Myara, I., Cosson, C., Moatti, N. & Lemonnier, A. Human kidney prolidase—purification, preincubation properties and immunological reactivity. Int. J. Biochem. 26, 207–214 (1994).

8. Jalving, R., Bron, P., Kester, H. C. M., Visser, J. & Schaap, P. J. Cloning of a prolidase gene from Aspergillus nidulans and characterisation of its product. Mol. Genet. Genomics MGG 267, 218–222 (2002).

9. Johnson, G. L. & Brown, J. L. Partial purification and characterization of two peptidases from Neurospora crassa. Biochim. Biophys. Acta 370, 530–540 (1974).

10. Kubota, Y., Shoji, S. & Motohara, K. Purification and properties of prolidase for germinating soybeans. Yakugaku Zasshi 97, 111–115 (1977).

11. Ghosh, M., Grunden, A. M., Dunn, D. M., Weiss, R. & Adams, M. W. Characterization of native and recombinant forms of an unusual cobalt-dependent proline dipeptidase (prolidase) from the hyperthermophilic archaeon Pyrococcus furiosus. J. Bacteriol. 180, 4781–4789 (1998).

12. Booth, M., Jennings, P. V., Nífhaolain, I. & O’cuinn, G. Endopeptidase activities of Streptococcus cremoris. Biochem. Soc. Trans. 18, 339–340 (1990).

13. Suga, K. et al. Prolidase from Xanthomonas maltophilia: Purification and Characterization of the Enzyme. Biosci. Biotechnol. Biochem. 59, 2087–2090 (1995).

14. Mikio, F., Yuko, N., Shigeyuki, I. & Toshio, S. Purification and Characterization of a Prolidase from Aureobacterium esteraromaticum. Biosci. Biotechnol. Biochem. 60, 1118–1122 (1996).

15. Fernández-Esplá, M. D., Martín-Hernández, M. C. & Fox, P. F. Purification and characterization of a prolidase from Lactobacillus casei subsp. casei IFPL 731. Appl. Environ. Microbiol. 63, 314–316 (1997).

16. Weaver, J., Watts, T., Li, P. & Rye, H. S. Structural Basis of Substrate Selectivity of E. coli Prolidase. PLOS ONE 9, e111531 (2014).

17. Wilk, P. et al. Substrate specificity and reaction mechanism of human prolidase. FEBS J. 284, 2870–2885 (2017).

18. Lupi, A. et al. Molecular characterisation of six patients with prolidase deficiency: identification of the first small duplication in the prolidase gene and of a mutation generating symptomatic and asymptomatic outcomes within the same family. J. Med. Genet. 43, e58 (2006).

19. Viglio, S. et al. The role of emerging techniques in the investigation of prolidase deficiency: from diagnosis to the development of a possible therapeutical approach. J. Chromatogr. B Analyt. Technol. Biomed. Life. Sci. 832, 1–8 (2006).

20. Phang, J. M., Liu, W. & Zabirnyk, O. Proline metabolism and microenvironmental stress. Annu. Rev. Nutr. 30, 441–463 (2010).

21. Wilk, P. et al. Structural Basis for Prolidase Deficiency Disease Mechanisms. FEBS J. (2018). doi:10.1111/febs.14620

22. Lupi, A., Tenni, R., Rossi, A., Cetta, G. & Forlino, A. Human prolidase and prolidase deficiency: an overview on the characterization of the enzyme involved in proline recycling and on the effects of its mutations. Amino Acids 35, 739–752 (2008).

23. Gecit, İ. et al. The Prolidase Activity, Oxidative Stress, and Nitric Oxide Levels of Bladder Tissues with or Without Tumor in Patients with Bladder Cancer. J. Membr. Biol. 250, 455–459 (2017).

24. Kucukdurmaz, F. et al. Evaluation of serum prolidase activity and oxidative stress markers in men with BPH and prostate cancer. BMC Urol. 17, (2017).

25. Surazynski, A., Miltyk, W., Palka, J. & Phang, J. M. Prolidase-dependent regulation of collagen biosynthesis. Amino Acids 35, 731–738 (2008).

26. Uygun Ilikhan, S. et al. Assessment of the correlation between serum prolidase and alpha-fetoprotein levels in patients with hepatocellular carcinoma. World J. Gastroenterol. WJG 21, 6999–7007 (2015).

27. Pirinççi, N. et al. Serum prolidase activity, oxidative stress, and antioxidant enzyme levels in patients with renal cell carcinoma. Toxicol. Ind. Health 32, 193–199 (2016).

28. Demir, S. et al. Decreased Prolidase Activity in Patients with Posttraumatic Stress Disorder. Psychiatry Investig. 13, 420–426 (2016).

29. Verma, A. K. et al. Association of Major Depression with Serum Prolidase Activity and Oxidative Stress. Br. J. Med. Med. Res. 20, 1–8 (2017).

30. Du, X., Tove, S., Kast-Hutcheson, K. & Grunden, A. M. Characterization of the dinuclear metal center of Pyrococcus furiosus prolidase by analysis of targeted mutants. FEBS Lett. 579, 6140–6146 (2005).

31. Phang, J. M., Pandhare, J. & Liu, Y. The Metabolism of Proline as Microenvironmental Stress Substrate. J. Nutr. 138, 2008S–2015S (2008).

32. Yang, L., Li, Y., Bhattacharya, A. & Zhang, Y. PEPD is a pivotal regulator of p53 tumor suppressor. Nat. Commun. 8, 2052 (2017).

33. Yang, L. et al. Prolidase directly binds and activates epidermal growth factor receptor and stimulates downstream signaling. J. Biol. Chem. 288, 2365–2375 (2013).

34. Yang, L., Li, Y. & Zhang, Y. Identification of prolidase as a high affinity ligand of the ErbB2 receptor and its regulation of ErbB2 signaling and cell growth. Cell Death Dis. 5, e1211 (2014).

35. Are, V. N. et al. Crystal structure of a novel prolidase from Deinococcus radiodurans identifies new subfamily of bacterial prolidases. Proteins Struct. Funct. Bioinforma. 85, 2239–2251 (2017).

36. Maher, M. J. et al. Structure of the Prolidase from Pyrococcus furiosus. Biochemistry 43, 2771–2783 (2004).

37. Bazan, J. F., Weaver, L. H., Roderick, S. L., Huber, R. & Matthews, B. W. Sequence and structure comparison suggest that methionine aminopeptidase, prolidase, aminopeptidase P, and creatinase share a common fold. Proc. Natl. Acad. Sci. U. S. A. 91, 2473–2477 (1994).

38. Lowther, W. T. & Matthews, B. W. Metalloaminopeptidases: common functional themes in disparate structural surroundings. Chem. Rev. 102, 4581–4608 (2002).

39. Lowther, W. T. & Matthews, B. W. Structure and function of the methionine aminopeptidases. Biochim. Biophys. Acta 1477, 157–167 (2000).

40. Lupi, A. et al. Human recombinant prolidase from eukaryotic and prokaryotic sources. Expression, purification, characterization and long-term stability studies. FEBS J. 273, 5466–5478 (2006).

41. Surażyński, A., Pałka, J. & Wołczyński, S. Phosphorylation of prolidase increases the enzyme activity. Mol. Cell. Biochem. 220, 95–101 (2001).

42. Surazynski, A., Liu, Y., Miltyk, W. & Phang, J. M. Nitric oxide regulates prolidase activity by serine/threonine phosphorylation. J. Cell. Biochem. 96, 1086–1094 (2005).

43. Lynch, M. & Marinov, G. K. The bioenergetic costs of a gene. Proc. Natl. Acad. Sci. U. S. A. 112, 15690–15695 (2015).

44. Bhatnager, R. & Dang, A. S. Comprehensive in-silico prediction of damage associated SNPs in Human Prolidase gene. Sci. Rep. 8, 9430 (2018).

45. Yoshimoto, T., Matsubara, F., Kawano, E. & Tsuru, D. Prolidase from bovine intestine: purification and characterization. J. Biochem. (Tokyo) 94, 1889–1896 (1983).

46. Falik-Zaccai, T. C. et al. A broad spectrum of developmental delay in a large cohort of prolidase deficiency patients demonstrates marked interfamilial and intrafamilial phenotypic variability. Am. J. Med. Genet. Part B Neuropsychiatr. Genet. Off. Publ. Int. Soc. Psychiatr. Genet. 153B, 46–56 (2010).

47. Ledoux, P., Scriver, C. & Hechtman, P. Four novel PEPD alleles causing prolidase deficiency. Am. J. Hum. Genet. 54, 1014–1021 (1994).

48. Cechowska-Pasko, M., Pałka, J. & Wojtukiewicz, M. Z. Enhanced prolidase activity and decreased collagen content in breast cancer tissue. Int. J. Exp. Pathol. 87, 289–296 (2006).

49. Wu, T.-J. et al. A framework for organizing cancer-related variations from existing databases, publications and NGS data using a High-performance Integrated Virtual Environment (HIVE). Database J. Biol. Databases Curation 2014, bau022 (2014).

50. Hornbeck, P. V. et al. PhosphoSitePlus, 2014: mutations, PTMs and recalibrations. Nucleic Acids Res. 43, D512–520 (2015).

51. Deribe, Y. L., Pawson, T. & Dikic, I. Post-translational modifications in signal integration. Nat. Struct. Mol. Biol. 17, 666–672 (2010).

52. Nussinov, R., Tsai, C.-J., Xin, F. & Radivojac, P. Allosteric post-translational modification codes. Trends Biochem. Sci. 37, 447–455 (2012).

53. Simão, F. A., Waterhouse, R. M., Ioannidis, P., Kriventseva, E. V. & Zdobnov, E. M. BUSCO: assessing genome assembly and annotation completeness with single-copy orthologs. Bioinforma. Oxf. Engl. 31, 3210–3212 (2015).

54. Yang, Y. et al. Dissecting Molecular Evolution in the Highly Diverse Plant Clade Caryophyllales Using Transcriptome Sequencing. Mol. Biol. Evol. 32, 2001–2014 (2015).

55. Rognes, T. Faster Smith-Waterman database searches with inter-sequence SIMD parallelisation. BMC Bioinformatics 12, 221 (2011).

56. Katoh, K. & Standley, D. M. MAFFT Multiple Sequence Alignment Software Version 7: Improvements in Performance and Usability. Mol. Biol. Evol. 30, 772–780 (2013).

57. Brown, J. W., Walker, J. F. & Smith, S. A. Phyx: phylogenetic tools for unix. Bioinformatics 33, 1886–1888 (2017).

58. Price, M. N., Dehal, P. S. & Arkin, A. P. FastTree 2 – Approximately Maximum-Likelihood Trees for Large Alignments. PLOS ONE 5, e9490 (2010).

59. Chatzou, M. et al. Generalized Bootstrap Supports for Phylogenetic Analyses of Protein Sequences Incorporating Alignment Uncertainty. Syst. Biol. (2018). doi:10.1093/sysbio/syx096

60. Sievers, F. et al. Fast, scalable generation of high-quality protein multiple sequence alignments using Clustal Omega. Mol. Syst. Biol. 7, 539–539 (2014).

61. Edgar, R. C. MUSCLE: multiple sequence alignment with high accuracy and high throughput. Nucleic Acids Res. 32, 1792–1797 (2004).

62. Blom, N., Gammeltoft, S. & Brunak, S. Sequence and structure-based prediction of eukaryotic protein phosphorylation sites. J. Mol. Biol. 294, 1351–1362 (1999).

63. Blom, N., Sicheritz-Pontén, T., Gupta, R., Gammeltoft, S. & Brunak, S. Prediction of post-translational glycosylation and phosphorylation of proteins from the amino acid sequence. Proteomics 4, 1633–1649 (2004).

64. Hunter, J. D. Matplotlib: A 2D Graphics Environment. Comput. Sci. Eng. 9, 90–95 (2007).

