## Supplementary figures and images for "Harnessing natural diversity to identify key amino acid residues in prolidase"

### Supplementary File 2

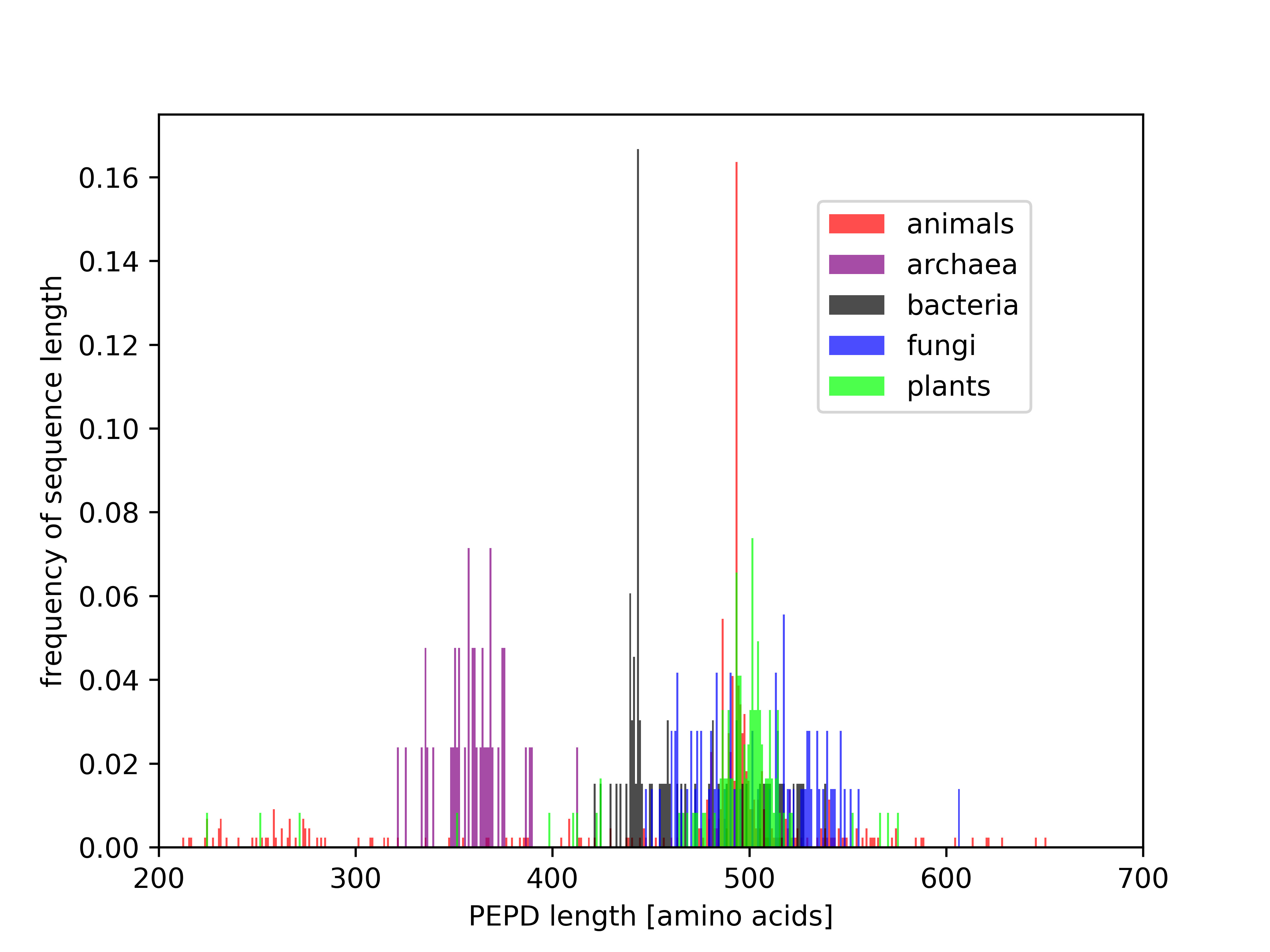

### Supplementary File 3

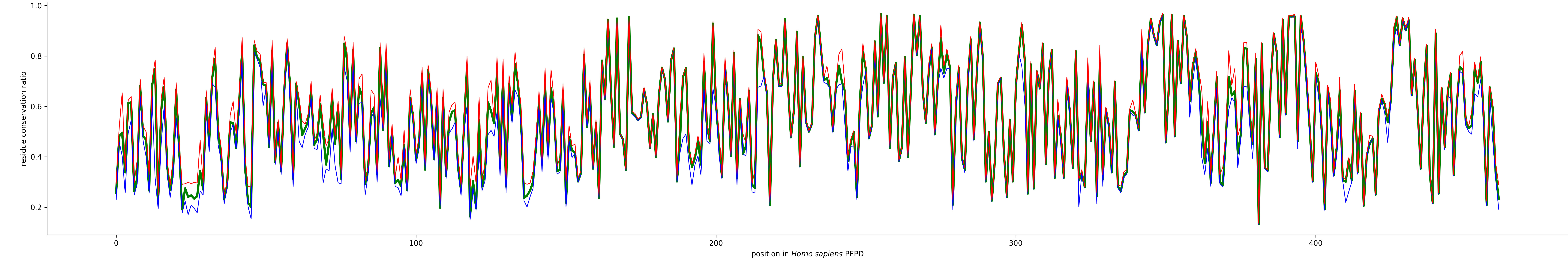

### Supplementary File 8

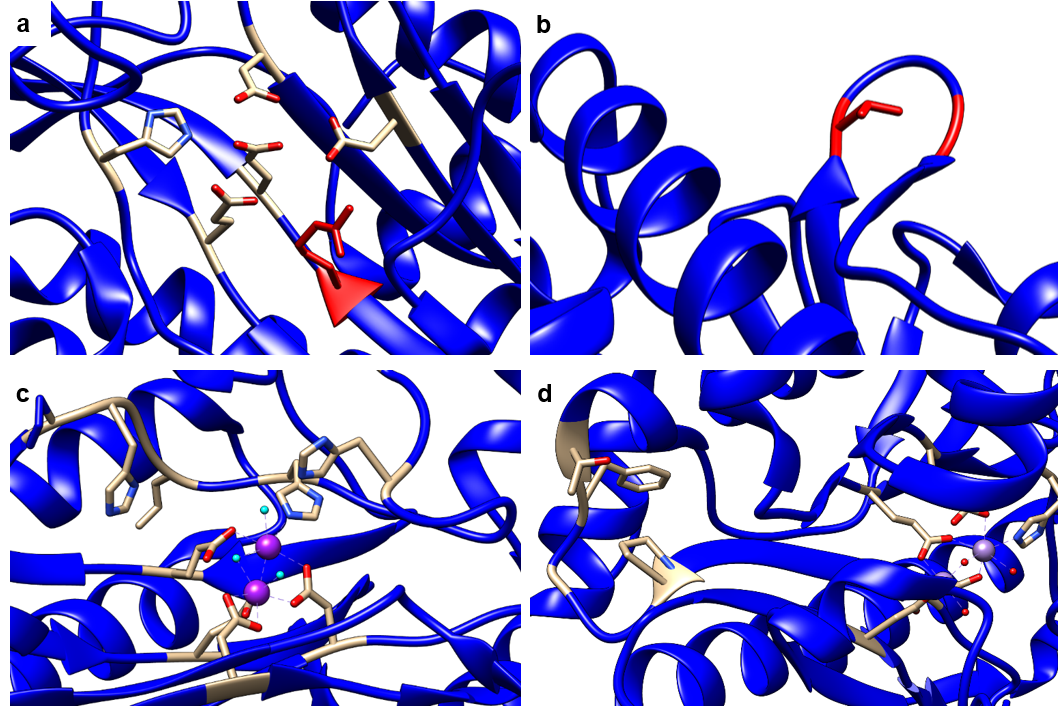

### Supplementary File 13

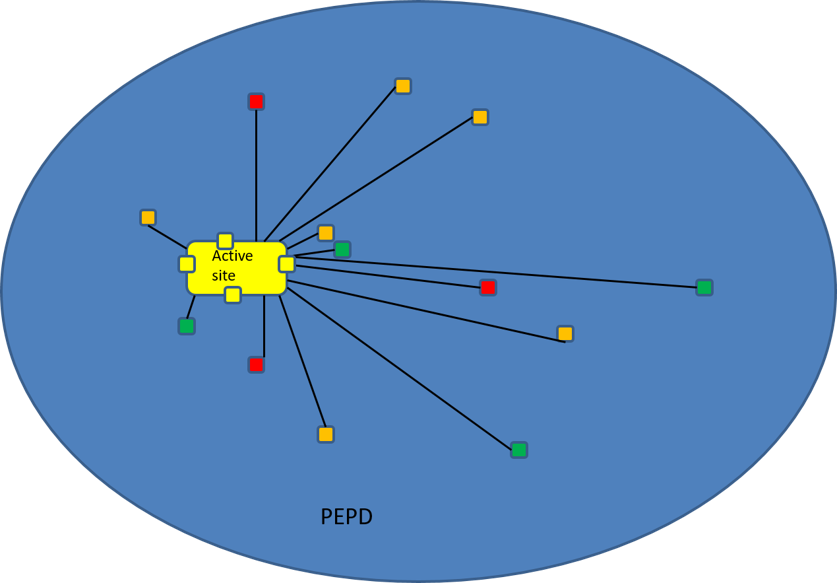
