## Supplementary File 10 for "Harnessing natural diversity to identify key amino acid residues in prolidase"

On average, eukaryotic data sets without detected PEPD homologs displayed about 15-20% lower completeness in the BUSCO analysis compare to data sets with PEPD candidates. This completeness difference for animals and plants was highly significant (p‑value << 0.01, Mann‑Whitney U test). Although fungi showed a substantial difference, it was not classified as statistical significant. There is no clear trend in archaea while bacteria display a high number of high quality data sets without PEPD candidates. This matches reports of *E. coli* PEPD mutants showing no observable phenotype thus supporting the assumption of a minor role in the majority of bacteria [1]. However, we validated that PEPD is present in 27 out of 32 analysed mycoplasma species. The mycoplasma data sets without a detected PEPD homolog showed on average 25% lower completeness compared to the data sets containing a PEPD candidate (SupplementaryFile 9). The completeness difference was highly significant (p‑value = 0.001, Mann Whitney U test). Therefore, technical reasons are likely to be one explanation for the lack of PEPD detection in these data sets. Due to the maximal reduced genome sizes of these species, they retain only the most important cellular functions, supporting the metabolic importance of PEPD [2].

In contrast, data sets of some species displayed multiple sequences with similarity to the human PEPD sequence even so *PEPD* seems to be a single copy gene in most species. The presence of one *PEPD* per haploid genome lead to multiple genes in polyploid species like *Triticum aestivum*. In addition, manual inspection revealed that fragmented gene models could be another explanation for this observation. Investigation of two *PEPD* copies in *Macaca mulatta* indicated the split of a gene into two predicted gene models. The annotated two genes are located on the same strand close to each other and encode different parts of PEPD (Macaca_mulatta@21290: 224 aa from putative N-terminus; Macaca_mulatta@3373: 258 aa from putative C‑terminus). Therefore, very short PEPD sequences were excluded from downstream analyses to prevent bias through partial sequences. However, not all sequences in the final data set have full length and are thereby affecting the maximal observed conservation that can be reached at a single position.

Further improvement of genomic sequences as well as additional RNA-Seq data sets might resolve these issues in the future by increasing assembly continuity and annotation quality. Theoretically, alternative transcripts of the same locus could pose another source of multiple peptide sequences per species. However, such cases are very rare as the annotation of most species is limited to one transcript and thus one resulting peptide sequence per locus.

**References**

[1]	Miller CG, Schwartz G. Peptidase-deficient mutants of Escherichia coli. J Bacteriol 1978;135:603–11.

[2]	Wilk P, Uehlein M, Kalms J, Dobbek H, Mueller U, Weiss MS. Substrate specificity and reaction mechanism of human prolidase. FEBS J 2017;284:2870–85. doi:10.1111/febs.14158.
